# Computational Insights Into Avidity of Polymeric Multivalent Binders

**DOI:** 10.1101/545012

**Authors:** E Zumbro, J Witten, A Alexander-Katz

## Abstract

Multivalent binding interactions are commonly found throughout biology to enhance weak monovalent binding such as between glycoligands and protein receptors. Designing multivalent polymers to bind to viruses and toxic proteins is a promising avenue for inhibiting their attachment and subsequent infection of cells. Several studies have focused on oligomeric multivalent inhibitors and how changing parameters such as ligand shape and size, and linker length and flexibility affect binding. However, experimental studies of how larger structural parameters of multivalent polymers such as degree of polymerization affect binding avidity to targets have mixed results with some finding an improvement with longer polymers and some finding no effect. Here, we use Brownian dynamics simulations to provide a theoretical understanding of how degree of polymerization affects the binding avidity of multivalent polymers. We show that longer polymers increase binding avidity to multivalent targets, but reach a limit in binding avidity at high degrees of polymerization. We also show that when interacting with multiple targets simultaneously, longer polymers are able to use inter-target interactions to promote clustering and improve binding efficiency. We expect our results to narrow the design space for optimizing the structure and effectiveness of multivalent inhibitors, as well as be useful to understand biological design strategies for multivalent binding.

## 1 Introduction

Biology uses multivalent interactions for a variety of reasons including enhancing weak monovalent interactions, creating conformal interfaces such as those between cells or those inducing endocytosis, or increasing specificity and affinity of binding using a limited number of receptor and ligand types [1]. Multivalent binding occurs when multiple ligands on one species bind to multiple receptors on another species simultaneously. While each individual binding site–ligand interaction might have weak binding affinity, when multiple sites bind simultaneously, they can produce a much stronger binding avidity than the sum of the corresponding monovalent interactions [1]. Here we use the term “avidity” as the overall binding affinity of multivalent interactions and “affinity” to refer to the binding affinity of single binding site interaction [2].

Because multivalent binding can be used to enhance low-affinity binding interactions such as those commonly found between glycoligands and sugar-binding proteins called lectins, designing synthetic multivalent polymers that target specific lectins is of great interest [3]. Binding strongly to lectins is a promising avenue for treating common diseases from diarrhea and colitis to influenza by inhibiting protein targets such as AB5 toxins including Shiga or Cholera toxin [4–9] or the hemaglutinin receptor on the influenza virus [10, 11].

To narrow the design space for multivalent inhibitors, several theoretical and experimental models have looked at how spacing of binding sites and flexibilities of linkers affect binding avidity [12–16]. Previous studies have explored optimizing parameters on the size scale of individual binding sites. For example, Liese et al. modelled how changing the linker length and flexibility between two ligands changes their binding avidity while Papp et al. explored matching the size between ligands exactly to the target [11, 17]. In contrast, the field has had relatively few studies on how large polyvalent materials and design decisions at size scales much larger than individual binding sites control polyvalent interactions. Some experimental studies have attempted to understand the effect of degree of polymerization on a polymer’s multivalent enhancement. Several of these studies found that longer polymers are more effective binders for influenza virus [10, 18, 19], but other groups have found that there is a limit to this binding enhancement from increased length when interacting with proteins [20, 21]. A general theoretical understanding of how polymer length contributes to multivalent binding has yet to be developed.

In this article, we show that while polymers with higher degrees of polymerization bind more tightly to multivalent targets, the enhancement in binding energy tapers off with polymer length. We find that the entropic penalty from forming long loops is a likely explanation for this effect. We also demonstrate that favorable interactions between targets, such as hydrophobic attraction between proteins, can enhance the binding of the inhibiting polymer to the target, but only for higher degrees of polymerization. We use a coarse grain Brownian dynamics simulation to establish rules for how the degree of polymerization can influence the strength of multivalent binding interactions between a polymer and a globular target such as a lectin.

## 2 Methods

We model our target as a single bead with multiple binding sites and our inhibitors as *N* freely jointed beads connected by harmonic springs as shown in Fig. 1. Each inhibitor bead has a single ligand. We model the chain using Brownian dynamics where the position of each polymer bead and target is governed by:

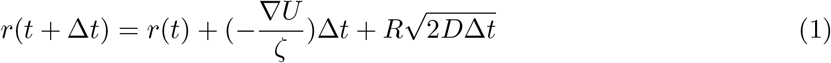

Where *r* is the position of the bead at time *t*, *R* is a random number drawn from a normal distribution with a mean of 0 and a standard deviation of 1, *ζ* is the drag coefficient, and *D* is the diffusion coefficient. The forces each bead experiences due to interactions with the surrounding polymer or toxin are captured in *∇U* where U is a potential energy that combines contributions from connectivity, excluded volume, and binding. These are added together as *U* = *U*_*sp*_ + *U*_*LJ*_ + *U*_*bind*_.

**Figure 1:**
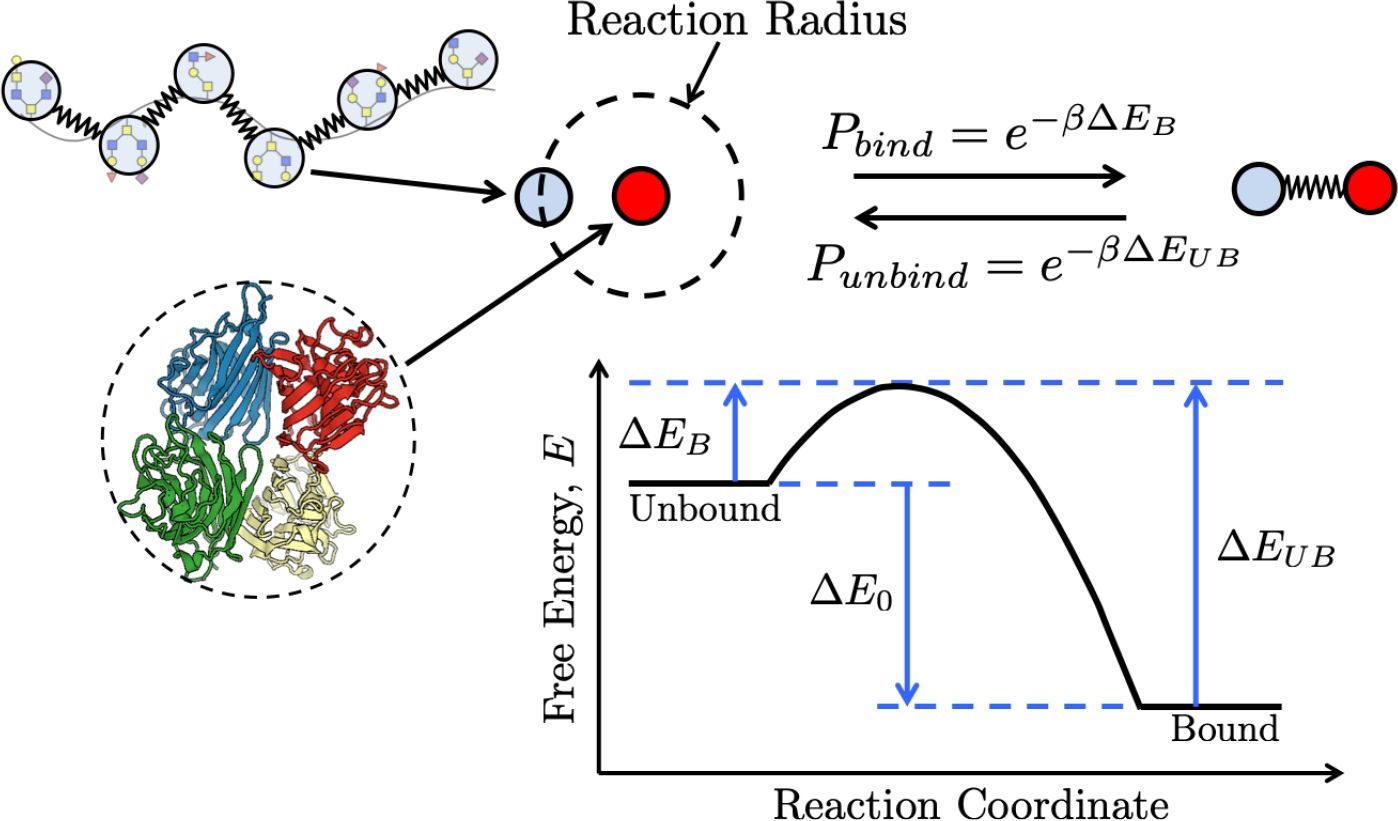
Inhibitors are represented by spherical beads (light blue) connected by gaussian springs. Each inhibitor bead has a single ligand. Targets can have multiple binding sites and are represented by a single spherical bead (red). Inhibitor ligands and target binding sites interact when they are within a reaction radius that is dependent on the timestep. Within this reaction radius, they have a probability of binding *P*_*bind*_ that depends on the depicted free energy landscape. Once bound, the target and inhibitor beads are connected by a gaussian spring, and with some probability *P*_*unbind*_ can return to being unbound and interacting solely through a Lennard-Jones potential. Rendering from the Protein Data Bank [22, 23].

The connectivity potential between adjacent polymer beads *U*_*sp*_ is modeled as a harmonic spring

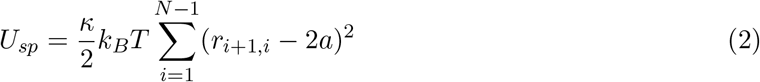

where *r*_*ij*_ is the distance between polymer beads, *a* is the radius of a simulation bead, and *κ* was chosen to be 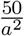, a value sufficiently large enough to prevent the polymer from stretching apart under normal Brownian forces.

A Lennard-Jones potential *U*_*LJ*_ was applied between bead pairs as

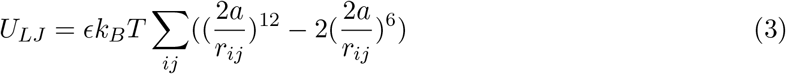

where the value of *ϵ* can be adjusted to control the solvent quality [24]. Across the simulations we used 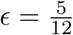 to mimic polymer configurations in a theta solvent and 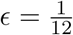 to mimic a good solvent as shown in Supplemental Figure S1. We chose to run both theta solvent and good solvent because these bound the solvent conditions for soluble polymers, putting a bound on any characteristics that depend on solvent quality.

To simulate reactive binding we apply a harmonic potential when two beads are bound and turn it on and off using a prefactor Ω(*i, j*).

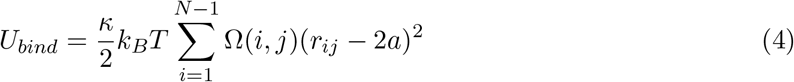

Ω(*i, j*) = 1 when the *i*th binding site on the target is bound to the *j*th bead of the inhibitor, and Ω(*i, j*) = 0 when the target binding site or inhibitor bead is unbound. To control the probability of binding and unbinding, we use a piecewise function based on the energy barriers for the binding reaction from C. Sing and A. Alexander-Katz [25].

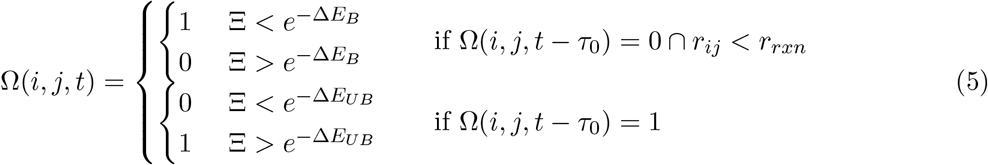

Here, Ξ is a random number between 0 and 1, ∆*E*_*B*_ is the energy barrier to bind normalized by *k*_*B*_*T*, and Δ*E*_*UB*_ is the energy barrier to unbind normalized by *k*_*B*_*T* as shown in Figure 1. Without loss of generality, these energies are considered to be always positive, and the kinetics of binding are held constant by keeping Δ*E*_*B*_ at 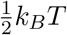 so that binding is an accessibly frequent event. The thermodynamic drive of binding is controlled by varying Δ*E*_0_ = Δ*E*_*B*_ − Δ*E*_*UB*_. Binding becomes more favorable as Δ*E*_0_ is made more and more negative. Binding reactions are evaluated every time interval *τ*_0_ = 100Δ*t*, where Δ*t* is the length of one timestep and *t* is the current time. The reaction radius *r*_*rxn*_ is the distance apart two beads would be if their surface was touching plus the distance that a bead can diffuse in *τ*_0_. Each inhibitor bead is only allowed to bind to one target binding site at a time. A target may have multiple binding sites.

The potentials are applied over the timestep 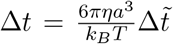 where 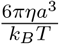 is the characteristic monomer diffusion time or the time that it takes a bead to diffuse its radius *a* and the dimensionless timestep is 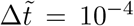. These equations can all be made dimensionless by scaling energies by thermal energy *k*_*B*_*T*, lengths by bead radius *a*, and times by the characteristic diffusion time 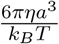.

## 3 Results and discussion

To better inform the design of multivalent polymeric binders, we seek to determine how degree of polymerization changes the inhibitor’s binding avidity. We examine two cases. In the first case, we observe how the binding avidity between a single monovalent or multivalent target and a polymer changes with the degree of polymerization of the polymer. In the second case, we probe how this result changes when the polymer is in the presence of multiple targets which have some favorable inter-target interactions.

### 3.1 Biologically relevant binding affinities

To establish a baseline binding affinity and ensure we are at biologically relevant binding affinities for individual binding sites, we first characterize the monovalent binding interactions of monovalent targets and monovalent free inhibitor beads. We place a single monovalent target in a box with a constant concentration of free inhibitor beads and measure the fraction of time the target is bound and unbound from an inhibitor bead. We run these simulations for 10^8^ timesteps and run either 50 or 100 simulations in parallel to ensure that we capture sufficiently long timescales that are much longer than the typical time for a single binding and unbinding to have accurate averaging. This is repeated at several different inhibitor bead concentrations. The resulting fraction of time bound plotted for different inhibitor concentrations is shown in Supplemental Figure S2. As expected, these lines fit the curve *θ* = [*I*]/([*I*] + *K*_*D*_), where *θ* is the fraction of time the target spends bound, [*I*] is the inhibitor concentration, and *K*_*D*_ is the dissociation constant. By looking at the *K*_*D*_, we ensure that we choose a Δ*E*_0_ of binding (Fig. 1) corresponding to an appropriate binding affinity for biologically relevant multivalent interactions. Assuming a target diameter of 5 nm, we find that using a Δ*E*_0_ = −5*k*_*B*_*T*, −4*k*_*B*_*T*, and − 2*k*_*B*_*T* corresponds to a *K*_*D*_ on the order of 4 × 10^−7^M, 1 × 10^−4^M, 8 × 10^−4^M respectively. This binding affinity is similar to the monovalent binding interactions between lectins and their corresponding sugars, which typically have a *K*_*D*_ in the mM to *μ*M range. [26, 27].

### 3.2 Effect of length on binding avidity to an individual toxin

Using Δ*E*_0_ = −4*k*_*B*_*T* as an experimentally relevant individual binding site affinity, we then explored how the degree of polymerization of our inhibiting polymer changed its binding avidity to a monovalent or multivalent target. We compared binding and unbinding kinetics of a single target with one or two binding sites to a constant concentration of inhibitor beads with varying connectivity. This scenario is depicted in Fig. 2, where a single target is placed with 64 inhibitor beads with increasing degrees of connectivity, such as 64 free inhibitor beads, 16 tetramers, or a single 64-mer.

**Figure 2:**
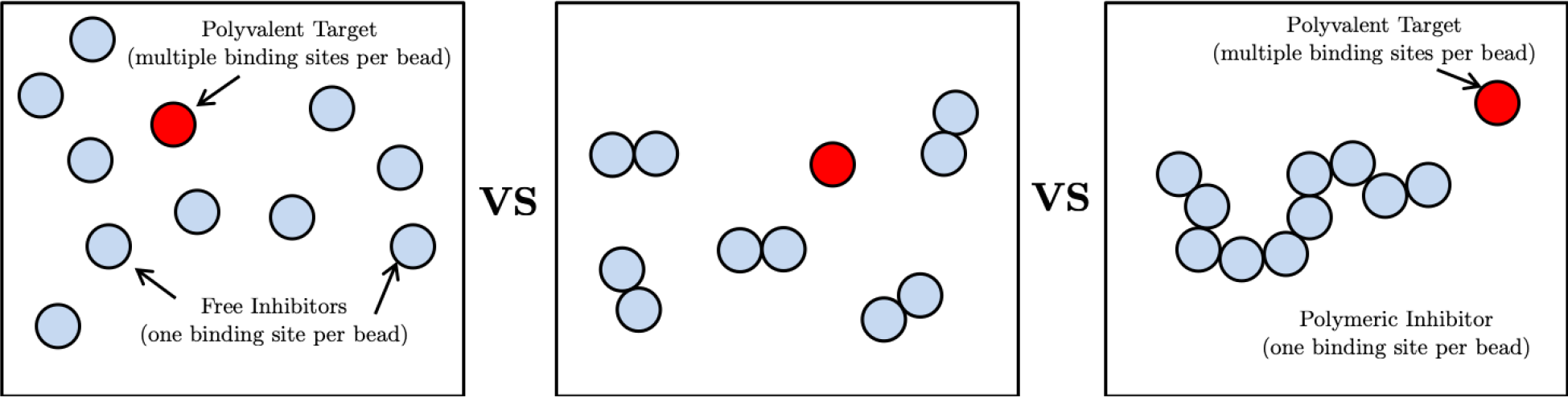
Schematic of the scenarios tested when comparing binding avidity’s dependence on degree of polymerization. The volume and number of inhibitor ligands are held constant to maintain a constant concentration of ligands at 64 ligands per box. The connectivity of the inhibitor ligands was varied from monomers to 64mers in multiples of two so that degrees of polymerization, 1, 2, 4, 8, 16, 32, and 64 were all investigated. This ensured that all polymers in each simulation were monodisperse.

To compare the binding avidity of the polymeric inhibitors to the target, we counted the time that a target stayed bound to an inhibitor bead, where a target was considered bound whenever at least one of the target’s binding sites was occupied. Unsurprisingly, the average time interval spent bound for monovalent targets does not depend on length (Fig. 3). The target’s single binding site can only interact with one inhibitor ligand at a time, so interactions with neighboring ligands have no impact on the duration the target spends bound. Therefore, the polymeric structure and valency of the inhibitor do not affect the average time bound for a single or dilute monovalent target.

**Figure 3:**
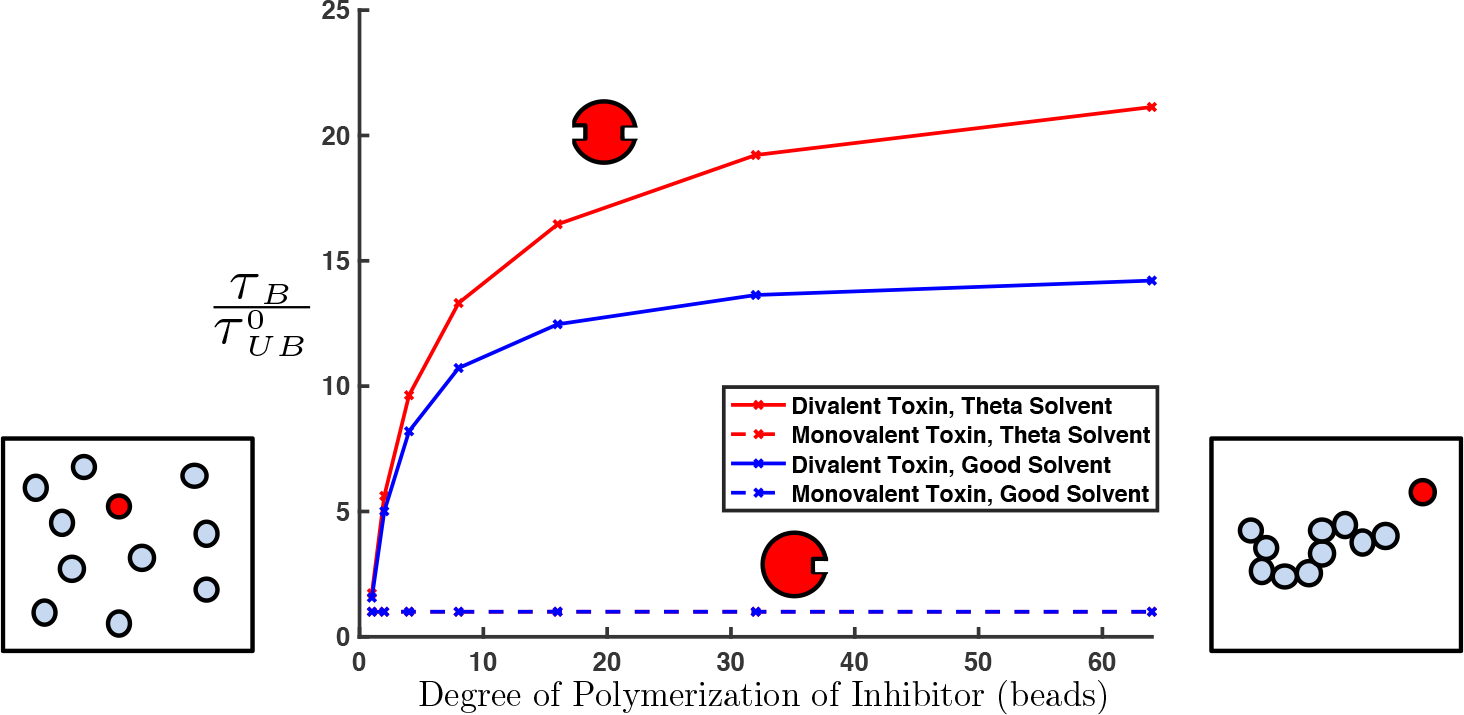
Average time interval a monovalent or divalent target are bound to polymeric inhibitors of various lengths, normalized by the average time bound for a monovalent target. The time interval bound for a monovalent toxin does not depend on the inhibitor length (dashed blue line and dashed red line overlap). For the divalent target, oligomeric inhibitors spend significantly more time bound than monomeric inhibitors, exhibiting the enhancement of multivalent binding avidities over monovalent binding. At high degrees of polymerization of the inhibiting polymer, as the length of the inhibitors is increased further, there is only a small gain in the average time interval the target is bound. The time spent bound approaches some maximum value with increasing inhibitor length.

In contrast, for the divalent targets, switching from monomeric inhibitors to polymeric inhibitors shows a significant increase in time the target spends bound (Fig. 3), demonstrating the enhancement in binding avidity created by multivalent binding. This follows the multivalent binding theories of increased local concentration [28], decreased loss of entropy over free ligands [1,2,29], and an increased rebinding effect [30]. Both the constant duration of time spent bound for our monovalent target and the increase in duration of time spent bound with the lengthening of our inhibiting polymer is consistent with all three of these previous theories.

However, these theories have not previously captured how degrees of polymerization much larger than the size scale of the target can change binding avidity. Revisiting Fig. 3, we can see that the time spent bound approaches a limit at high degrees of polymerization for both theta 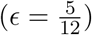 and good solvent qualities 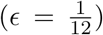. To explain this phenomenon, we considered the proportion of time a target is bound in a system with a given degree of polymerization. This proportion can be transformed into a free energy of binding, which we term Δ*G*_*B*_:

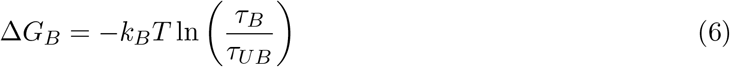

where *τ*_*B*_ and *τ*_*UB*_ are the average time spent bound and unbound respectively. While *τ*_*B*_ as examined in Fig. 3 is difficult to treat theoretically, we found this Δ*G*_*B*_ more theoretically tractable. We developed a model predicting Δ*G*_*B*_ as a function of the degree of polymerization and valency of the target as well as other factors, described in detail in Supplemental Information. Briefly, the model is loosely inspired by the Poland-Scheraga model of DNA denaturation, in that a polymer bound multivalently to a target can be represented as a sequence of loops alternating with sites bound to the target [31]. In the general case, the partition function of this model can only be evaluated numerically, but in the limit of high *N*_*P*_, where *N*_*P*_ is the degree of polymerization, there is an analytical result for Δ*G*_*B*_. The full function, given in Eq. S21, is complex, but the dependence on *N*_*P*_, the number of polymers *n*, and volume *V*_*box*_, is simple:

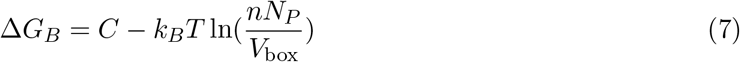

where *C* is a value not dependent on *N*_*P*_ related to the persistence length, solvent quality, ligand density, and valency of the target. Note that in our simulations, both *nN*_*P*_ and *V*_*box*_ are held constant; specifically, *nN*_*P*_ = 64. Thus, Eq. 7 predicts that at high *N*_*P*_, polyvalency no longer increases avidity. So, for example, if *N*_*P*_ = 32 is high enough to approach the limit (a question we address shortly), Δ*G*_*B*_ should be the same for two 32-mers and one 64-mer. Our theoretical treatment successfully reproduces the qualitative behavior of Δ*G*_*B*_: as predicted, we observe that Δ*G*_*B*_ initially decreases sharply, representing the benefits of polyvalency, before reaching a limit at higher degrees of polymerization, shown in Fig. 4.

**Figure 4:**
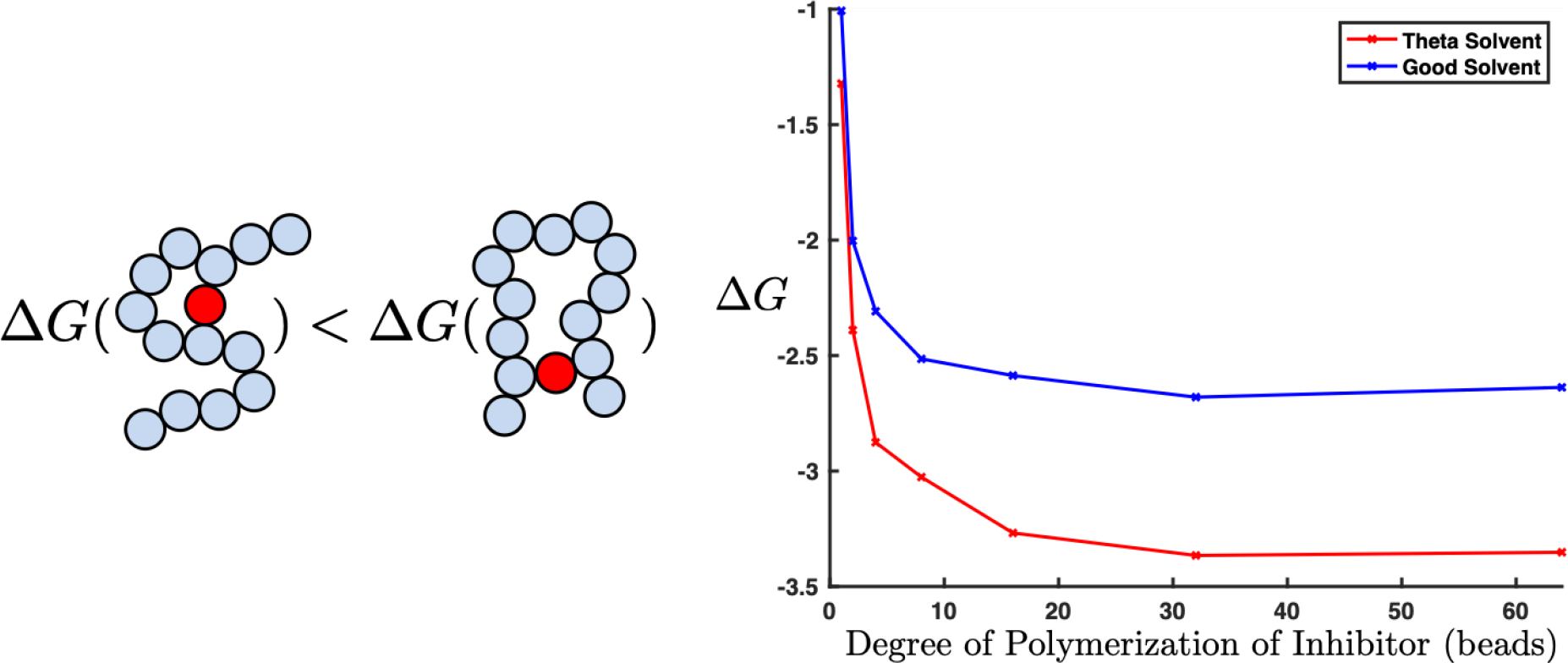
A plot of the free energy of binding for the degree of polymerization of the inhibitor. The free energy of binding is calculated using the average time interval the target spends bound to the polymer (meaning one or more binding sites is bound) divided by the average time interval the target spends completely unbound. Longer loops are entropically unfavorable, so while they are possible in longer polymers, they are unlikely to form. We can see that this leads to a limiting minimum binding energy as degree of polymerization of our inhibitor increases. This is true in both good (blue) and theta (red) solvents.

The leveling off of Δ*G*_*B*_ in our theoretical model is due to loop entropy: when two faraway monomers each bind the target, the polymer is forced into a large loop, which restricts the conformational degrees of freedom of the polymer chain. This free energy penalty increases with the size of the loop, and for large enough loops, the free energy penalty becomes larger than the free energy of binding. Beyond the length where binding loops are no longer thermodynamically favored (which corresponds to where the high-*N*_*P*_ limit begins to be reached), increasing the degree of polymerization will provide no benefit. Thus, we predicted that the flattening of the Δ*G*_*B*_ curve should coincide with the length at which loops stop forming. Indeed, the frequency of loops drops precipitously with loop lengths, and loops larger than a length of 9 for good solvent and 13 for theta solvent are vanishingly rare as shown in Fig. 5. This is in agreement with the fact that Δ*G*_*B*_ flattens beyond *N*_*P*_ = 8 for good solvent and *N*_*P*_ = 16 for theta solvent (Fig. 4). Thus, loop entropy is the likely culprit for the diminishing returns of increasing *N*_*P*_.

**Figure 5:**
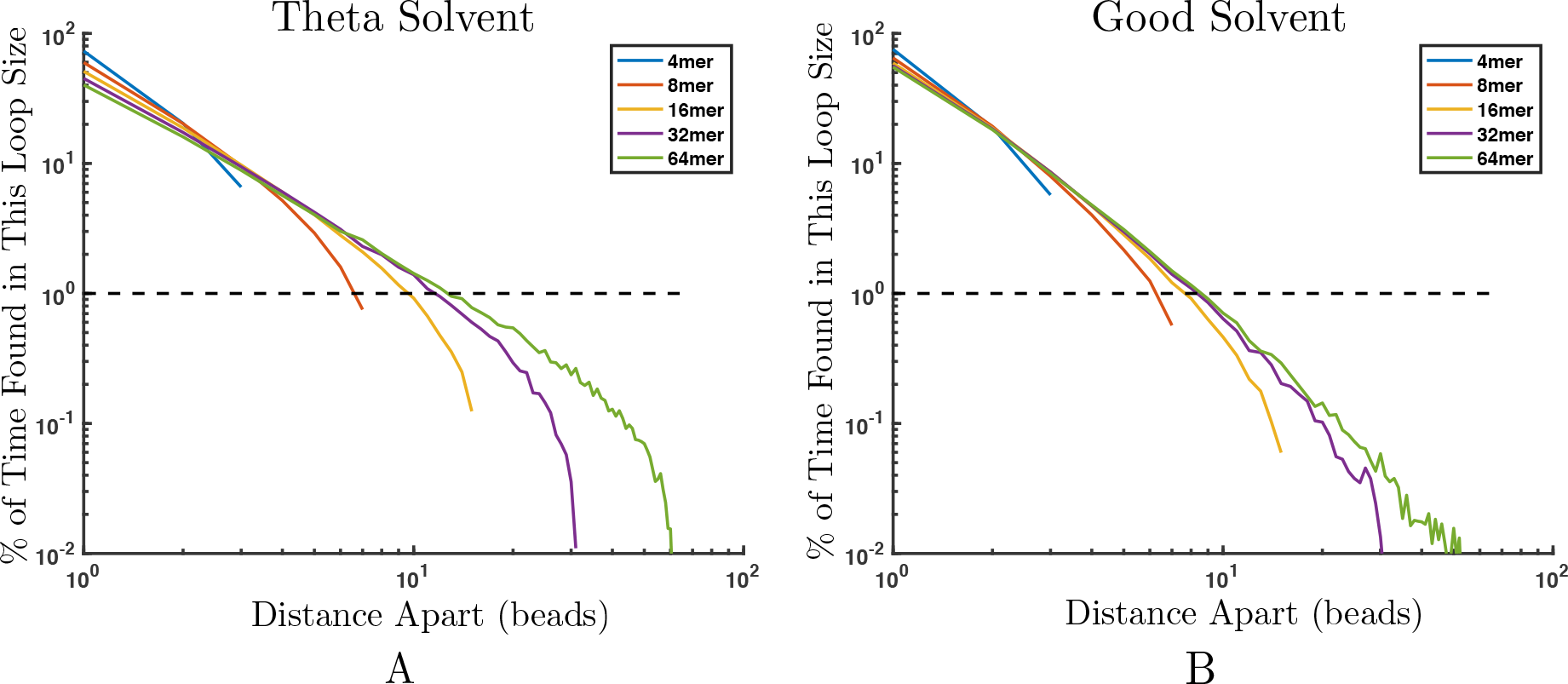
Log-log plot of the percent of time a divalent target forms various loop sizes for different length polymers. For reference, 1% frequency is shown with dashed black line. In theta solvent (A) and good solvent (B), loops larger than 13 and 9 beads, respectively, are formed less than 1% of the time.

Relatedly, we predict that solvent quality should not only affect the polymer length at which diminishing returns on avidity are reached; specifically, the maximum avidity should be weaker in good solvent due to the increased entropic cost of forming large loops. This prediction is also in agreement with the results from Fig. 4. Ultimately, our simulation and model results match excitingly well with the experimental results that increasing polymer length leads to only a limited increase in polymer avidity to lectins [20, 21].

### 3.3 Effect of length on binding avidity in the presence of multiple toxins

In vivo, environments can be crowded and multiple targets might interact with a single inhibiting polymer. If the target is a protein, hydrophobicity and charge can create target-target interactions leading to a wide range of solubilitiy maximums from 1 mg/ml for wheat germ agglutinin to more than 50 mg/ml for serum albumin [32, 33]. In this section, we examine binding between multiple targets and the inhibiting polymer and consider how target-target interactions influence binding avidity. To investigate the affect that target-target interactions have on the binding avidity of the inhibitors, we added a Lennard-Jones potential between targets and explored how changing the attraction between the targets modified their binding with the inhibiting polymer.

To examine the effect of multiple targets interacting with the inhibitor simultaneously, we placed 64 divalent targets in a box with inhibiting polymers. To compare the effect of polymer length, we again varied the connectivity of the inhibiting beads while maintaining the same total concentration of polymer binding sites, as depicted in Fig. 6. We modified our target-target attraction by changing *E* in Eq. 3, and compared two target-target attraction scenarios: a relatively neutral condition where 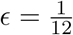 and a weakly attractive condition where 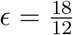. To ensure we were at biologically relevant target-target interactions, we calculated the concentration of our targets by making the following assumptions. Assuming a target diameter of 5 nm and molecular weight of 70 kDa, 64 targets corresponds to a concentration of 7 mg/ml. By running 64 targets in a box without an inhibiting polymer present, we confirmed that at both 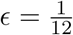 and 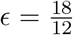 the targets are not attracted enough to aggregate on their own. This shows that both levels of target-target Lennard-Jones interactions are within the range of relevant protein solubilities.

**Figure 6:**
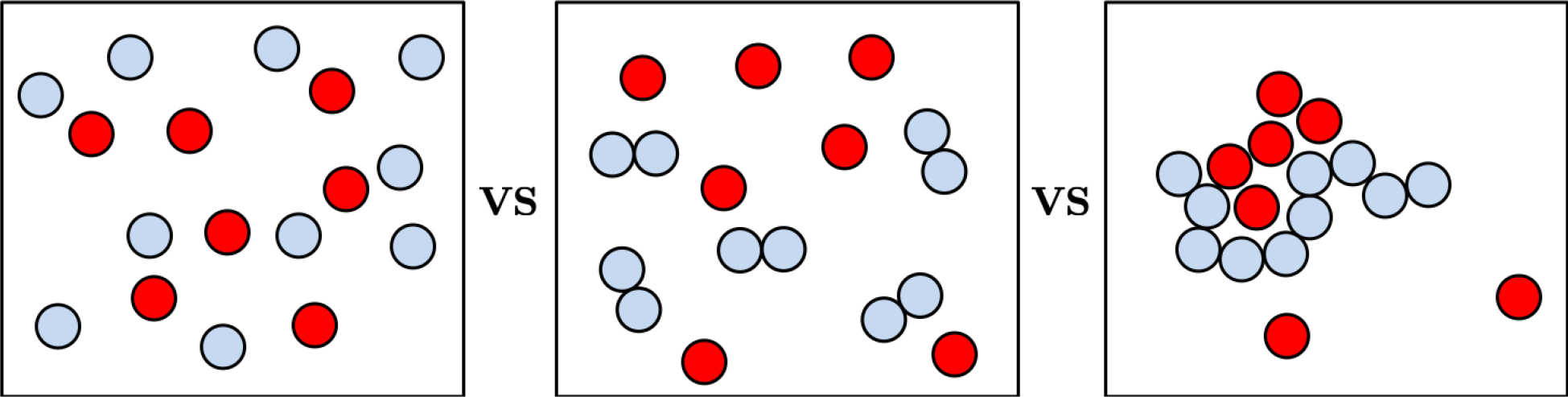
Schematic of the scenarios tested when comparing binding avidity’s dependence on degree of polymerization with multiple targets present. The volume and number of inhibitor ligands are held constant to maintain a constant concentration of ligands at 64 ligands per box. The connectivity of the inhibitor ligands was varied from monomers to 64mers in multiples of two so that degrees of polymerization, 1, 2, 4, 8, 16, 32, and 64 were all investigated. This ensured that all polymers in each simulation were monodisperse. The concentration of targets was held constant in all simulations.

#### 3.3.1 Increased competition

Normally one does not have isolated targets, but a finite concentration of them. Thus, it is interesting to ask the following question: if one had multiple targets with a given degree of solubility binding to the same inhibitor, would that have a marked effect on the kinetics? To answer this, we examined the binding kinetics of 64 targets to our inhibiting polymers to compare to our single or dilute target case. Similarly to when interacting with single targets, the binding avidity of polymers initially increases with increasing degree of polymerization before tapering off at high polymerization as shown in Fig. 7. More interestingly, in the presence of multiple targets, increased attraction between targets decreases the maximum *τ*_*B*_. To investigate this phenomenon, we compared the rate of unbinding in Fig. 8. Here, we see two timescales at which targets unbind, a fast and a slow timescale. The fast timescale represents targets that only become singly bound before unbinding, whereas the slow timescale represents targets that transition from being doubly bound to unbound. By comparing the slope of the linear best fit line in both regions, we find that the rate of unbinding for single bonds is unchanged when there is inter-target attraction, but the rate of unbinding for doubly bound targets increases with inter-target attraction. The increased rate of unbinding for doubly bound targets leads to the decrease in average *τ*_*B*_ seen in Fig. 7.

**Figure 7:**
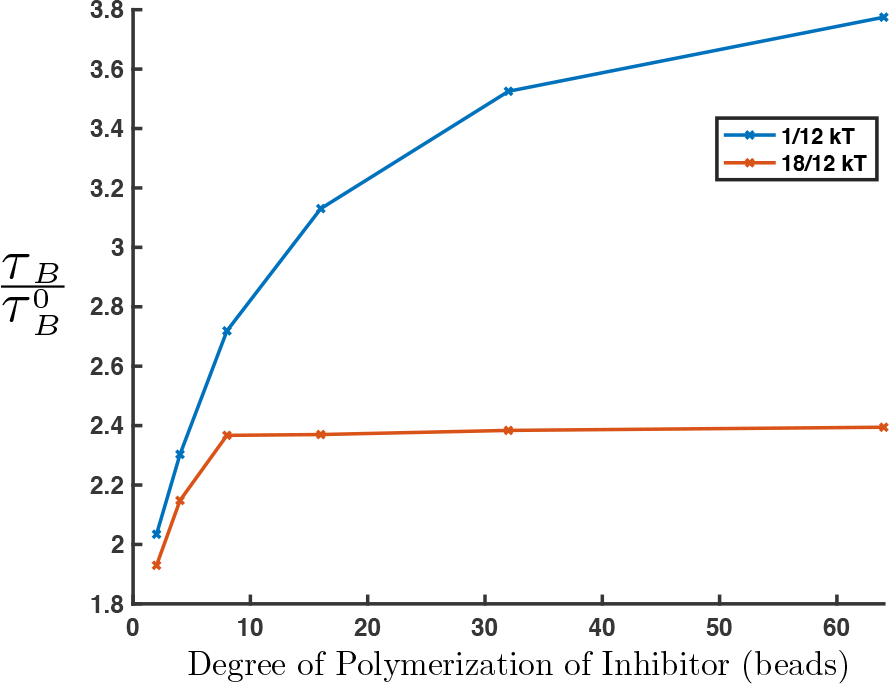
Plot of average time bound (*τ*_*B*_) for targets when multiple targets are present. Y-axis is normalized by the average time bound for monomeric inhibitors, 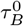. Data presented is for polymer-target binding energy of Δ*E*_0_ = −4 *k*_*B*_*T* in theta solvent. Similar to with a single target, *τ*_*B*_ has a limited increase with degree of polymerization of the inhibiting polymer. More attractive inter-target potentials (orange) decrease the maximum *τ*_*B*_.

**Figure 8:**
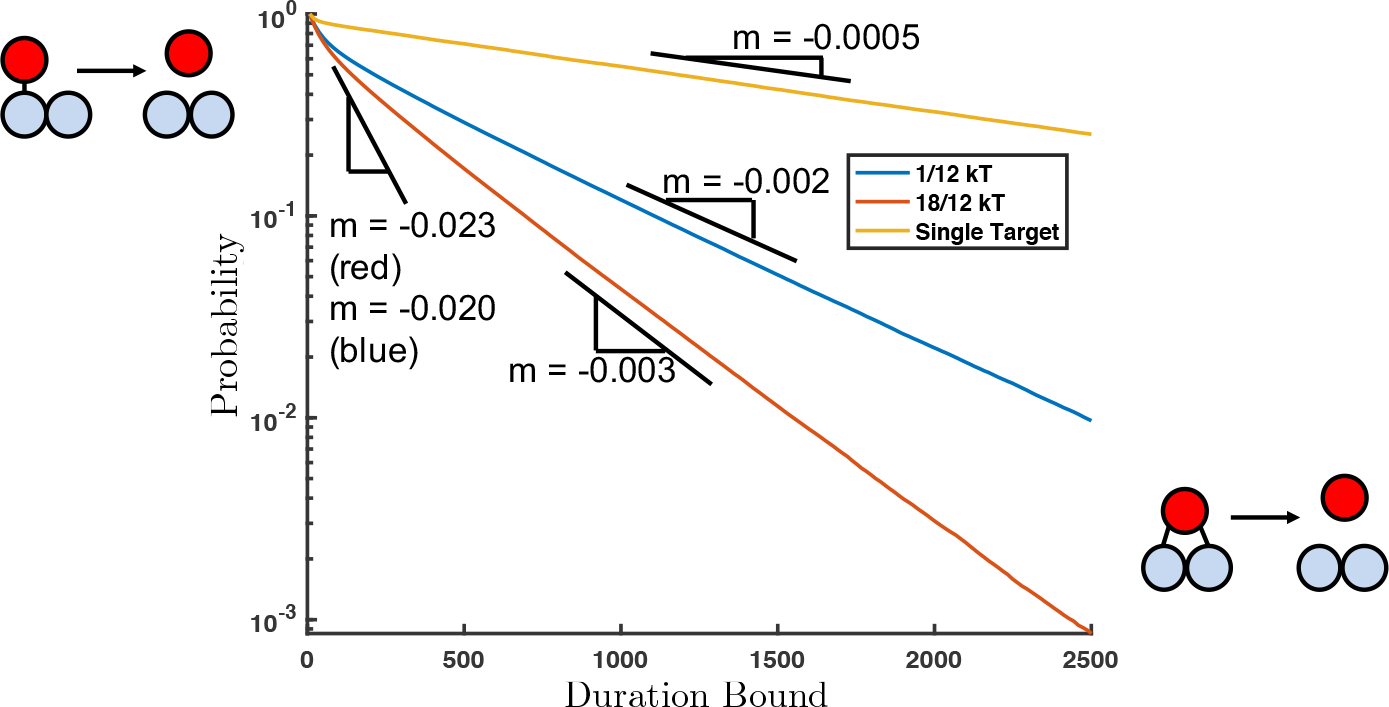
Distribution of times spent bound for single target scenarios (yellow) and scenarios with multiple targets (orange and blue). The rate of unbinding corresponds to the slope of the line in these two regions, shown in black. There is a fast and a slow timescale on which targets unbind. The former corresponds to singly bound targets unbinding and does not change with inter-target potentials. The second, longer timescale corresponds to doubly bound targets that unbind. When there are favorable target-target interactions (orange), the decay in doubly bound times is slower than if there is not an attraction between targets (blue), or if there is no competition from other targets (yellow).

The higher probability that doubly bound targets unbind can be explained by increased competition. If a lone or very dilute target becomes doubly bound and then unbinds with one binding site, this unbound site could easily rebind. In contrast, in a crowded environment with many targets, a site that unbinds has to compete with neighboring targets to rebind. This increase in competition comes from both neighboring bound targets (Fig. 9A) as well as unbound targets that are aggregated by a high density of bound targets (Fig. 9B). We will show that the increase in inter-target attraction from 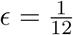 to 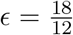 leads to a drastically higher local concentration of targets in the polymer’s radius of influence, exacerbating this competition and shortening the maximum *τ*_*B*_.

**Figure 9:**
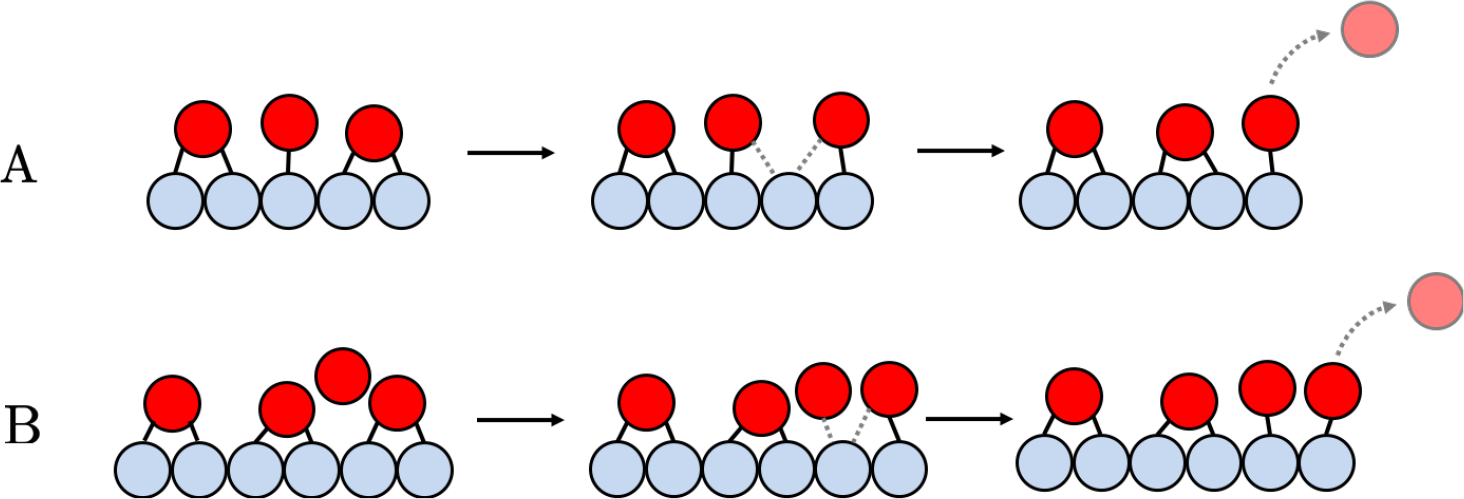
There are two types of competition bound targets experience that lead to shorter times bound for divalently bound targets. (A) shows competition from neighboring bound targets and (B) shows competition from nearby unbound targets which is increased for more favorable inter-target potentials.

#### 3.3.2 Polymer induced phase separation

Because the kinetic changes were correlated to changes in local concentration of the target, we next considered the thermodynamics of the system, where we found an increased concentration of targets bound to the inhibitor. In Fig. 10A, we show that in theta solvent for ~ 0.1 mM binding affinity (Δ*E*_0_ = −4*k*_*B*_*T*), the average number of targets bound to the polymer increased for higher target-target attraction, for both mono and divalent targets. Therefore, although individual targets unbind more quickly, inter-toxin attraction leads to higher inhibiting polymer avidity overall.

**Figure 10:**
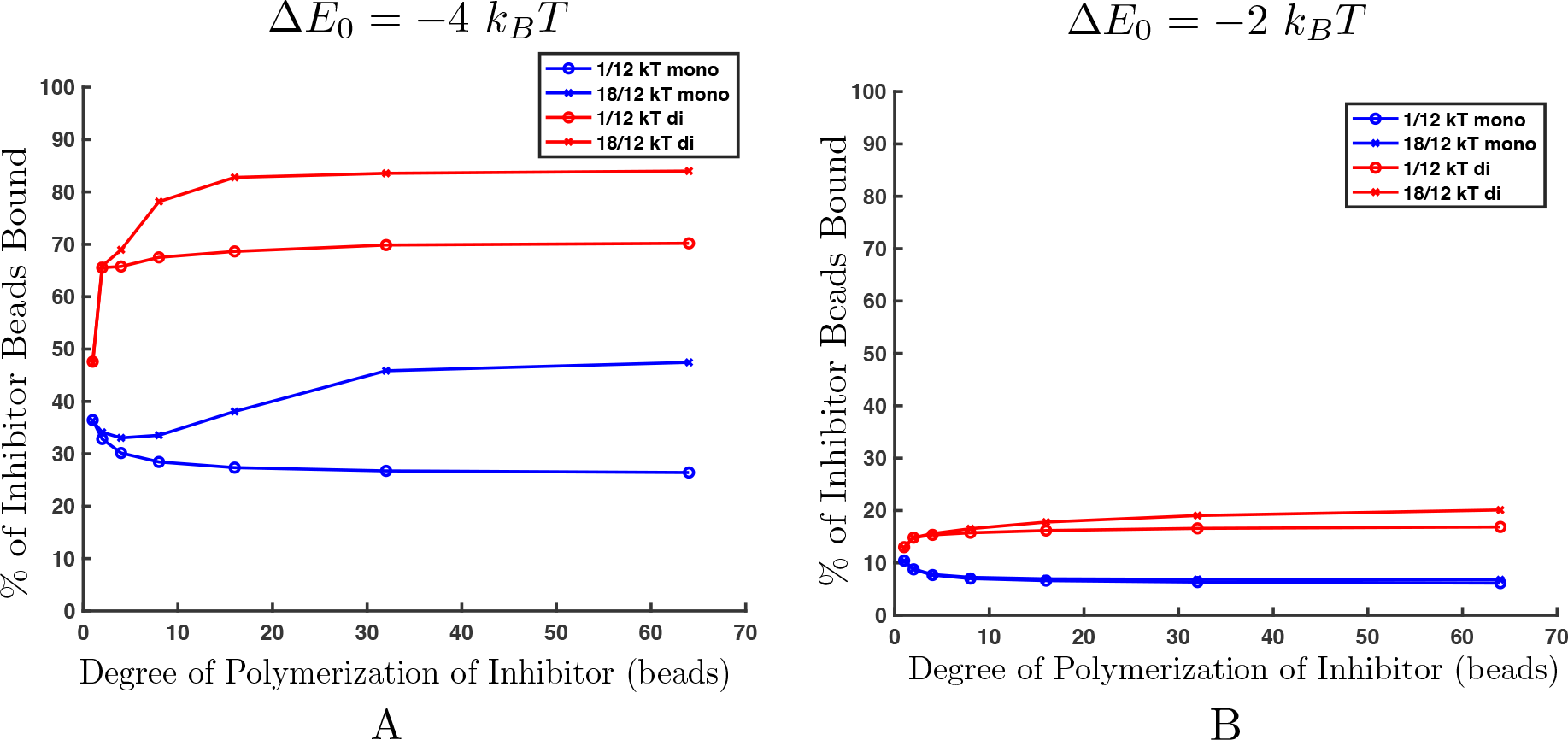
Percent of inhibitor beads bound in theta solvent when the target-polymer binding affinity is Δ*E*_0_ = −4 *k*_*B*_*T* (A) and −2 *k*_*B*_*T* (B). (A) As inhibitor length increases, a transition occurs that allows the polymer to bind a significantly higher percentage of targets when there is some inter-target attraction. This transition happens at approximately degree of polymerization of 10 for monovalent targets and degree of polymerization of around 3 for divalent targets. (B) At very low polymer-target binding affinities, such as −2 *k*_*B*_*T*, a critical percentage of targets never bind to the inhibiting polymer, so even at high degrees of polymerization, a transition in binding avidity does not occur.

Attraction between targets causes a significant increase in the number of targets bound because it induces a collapse transition where bound targets collapse the polymer and themselves into a globule or liquid phase. When the polymer/bound target system collapses to form a globule, the enthalpic benefit of an additional target joining the globule becomes greater than the loss of entropy of binding, leading to a significant increase in the number of targets bound to the target. This leads to a target rich liquid-like phase attached to the polymer and a low concentration gas-like target phase in the supernatant. Similar data for good solvent can be seen in Supplemental Figure S3.

We confirmed that the marked increase in avidity was caused by a polymer collapse transition by examining the end to end distance in Fig. 11. Fig. 11 shows the decrease in the average end to end distance for a 64mer polymer in theta solvent interacting with divalent (Fig. 11A) and monovalent (Fig. 11B) targets with 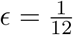 and 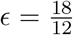 attractions between targets. End to end distances for polymers in good solvent interacting with multiple targets can be seen in Supplemental Figure S4. As expected, the theta polymer is at its normal random walk size of 8 with no targets present, but when divalent targets are added, the polymer collapses to a globule for both levels of inter-target attraction. For 64mers interacting with monovalent targets, we only see a collapse in the end to end distance when the inter-target attraction is 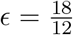, meaning that the collapse transition does not occur when 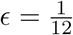. Indeed, this is reflected in our plot in Fig. 10A of the average number of monovalent targets bound. For 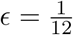, long polymers bind fewer monovalent targets than the monomeric inhibitors due to the high translational entropy cost of targets binding to a large polymer.

**Figure 11:**
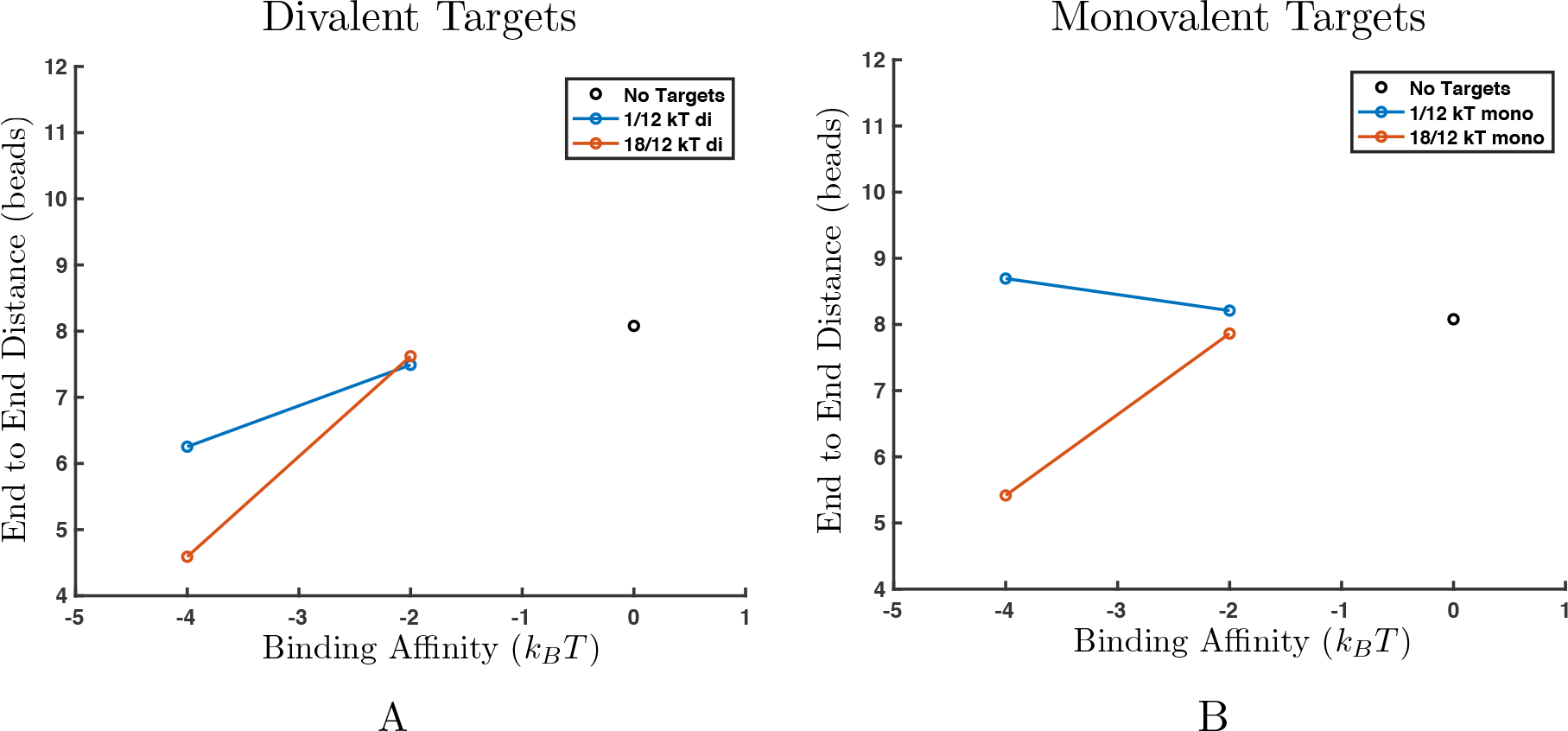
End to end distance for 64-mer polymers in theta solvent in the presence of divalent targets (A) and monovalent targets (B). (A) Increasing binding affinity between the targets and polymers induces a transition where the polymer collapses in size for both inter-target attractions. (B) Only high inter-target attraction leads to a collapse transition (orange). Low inter-target attraction (blue) does not provide enough enthalpic gain to overcome the entropic loss of phase separation.

This collapse transition that leads to globular polymers and higher target binding is caused by a competition between entropy and enthalpy and can be induced by increasing the polymer length. This can be seen in the large jump in targets bound for monovalent targets with 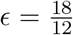 in Fig. 10A as the degree of polymerization is increased from 8 to 32 beads. Examining this case more closely, we see that as degree of polymerization is increased, the percent of inhibitor beads bound initially drops, before a sudden increase in binding after a polymerization of approximately 10 beads. At low degrees of polymerization, translational entropy of the targets dominates the binding interaction, so the targets prefer to bind to inhibitor beads with higher translational degrees of freedom. If there is low inter-target attraction, targets bind most favorably to monomeric inhibitors because monomeric inhibitors have the highest degrees of freedom. Consequently, fewer targets bind to longer polymers because they have fewer ends and the binding sites in the middle of the polymer have fewer translational degrees of freedom. This decrease in binding affinity with increasing polymer length is shown nicely in in Fig. 10A for the monovalent targets when 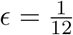. We further confirm this in Fig. 12A, where we see that monomeric inhibitors spend a relatively uniform amount of time bound, whereas the ends of a 64mer spend almost 50% more time bound than the center beads and overall spend less time bound than the monomeric inhibitors.

**Figure 12:**
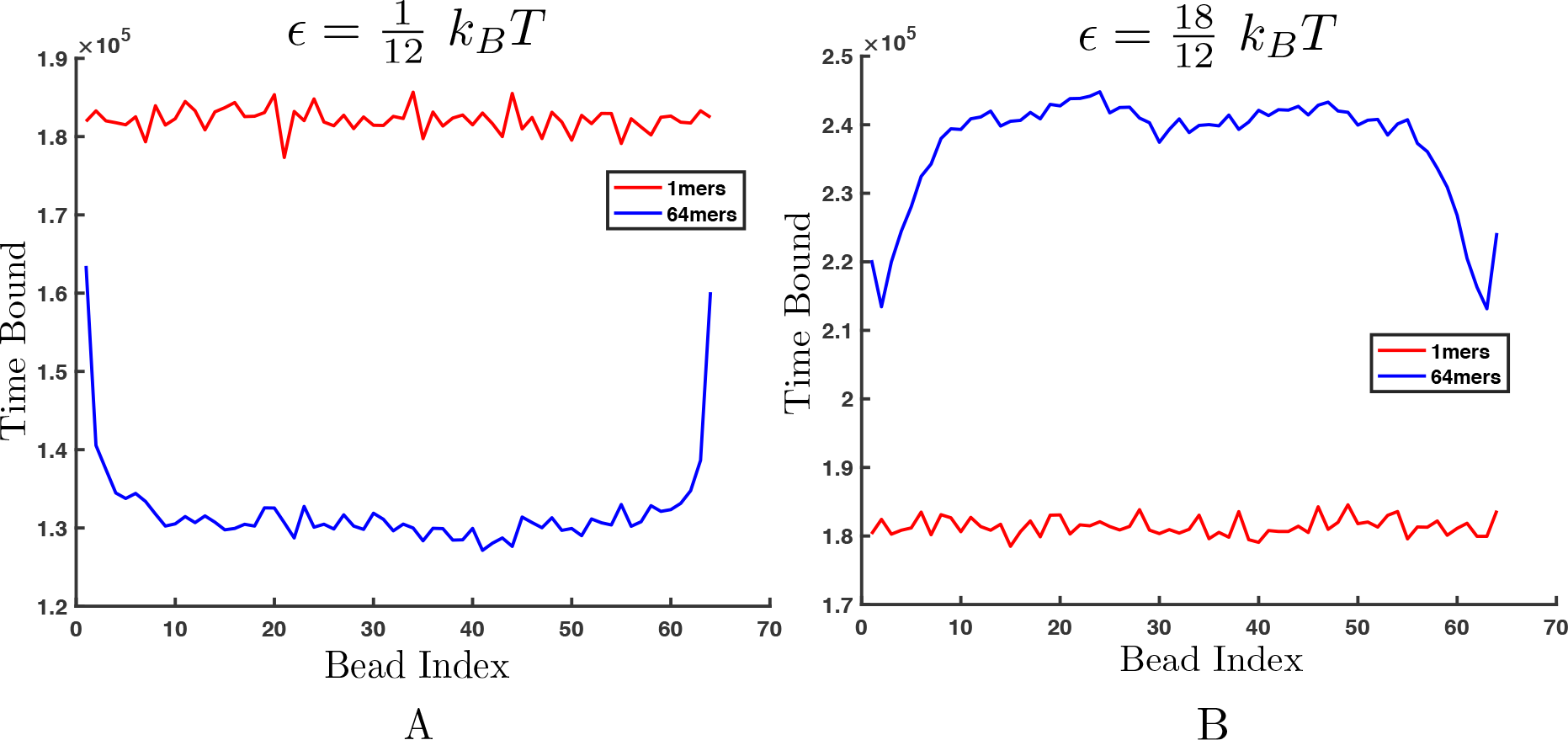
The amount of time each inhibitor bead spends bound when interacting with monovalent, −4 *k*_*B*_*T* binding targets. Plots compare binding times for monomeric inhibitor beads (red) and beads that are part of a 64-mer polymer (blue). (A) Time bound when interacting with targets that have low (*ϵ* = 1/12 *k_B_T*) target-target attraction. Monomers are each bound for a uniform amount of time, but the polymer ends are bound much more frequently than the polymer beads in the center of the chain. (B) When interacting with targets that have higher target-target attraction (*ϵ* = 18/12 *k_B_T*), the polymer collapses, making the center beads bound more frequently than the chain ends. Monomeric inhibitor beads continue to experience uniform binding preference.

Therefore, in Fig. 10A, when there is initially a decrease in the monovalent targets bound for 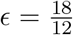, we are seeing the domination of the translational entropy term at low degrees of polymerization. At high degrees of polymerization, enthalpy starts to dominate and the polymer collapses, as shown in Fig. 11. Examining which polymer beads spend more time bound in Fig. 12B, we see that for 64mers, a significant transition has occurred where the center beads spend considerably more time bound than the ends or monomeric inhibitors. This can be explained by the combined enthalpic benefit of binding to the polymer and the favorable enthalpy of being nearby other targets. The ends of the polymer remain bound less frequently because targets that bind at the polymer ends have on average fewer target neighbors and thus, binding to ends has less enthalpic benefit.

In addition to increased binding of targets, the polymeric inhibitor also promotes aggregation and increased local concentration of unbound targets. By measuring the minimum distance between all unbound targets and the polymer and normalizing by the volume of the shell, we compared the concentration of targets at each distance *R* away from the polymer as shown in Fig. 13 for theta solvent and Supplemental Figure S5 for good solvent. From these plots, it is clear that at small inter-target potentials such as 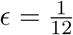 there is a negligible increase in the concentration of unbound targets near the polymer. In contrast, with an attractive inter-target potential of 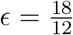, there is a significant increase in the concentration of unbound targets near the polymer - almost 5 times the bulk concentration for theta solvent and Δ*E*_0_ = −4*k*_*B*_*T*. Overall, this means that intertarget attraction leads to significant increases in both bound targets and unbound target clustering, encouraging the collapse transition that makes the polymeric inhibitors more effective binders.

**Figure 13:**
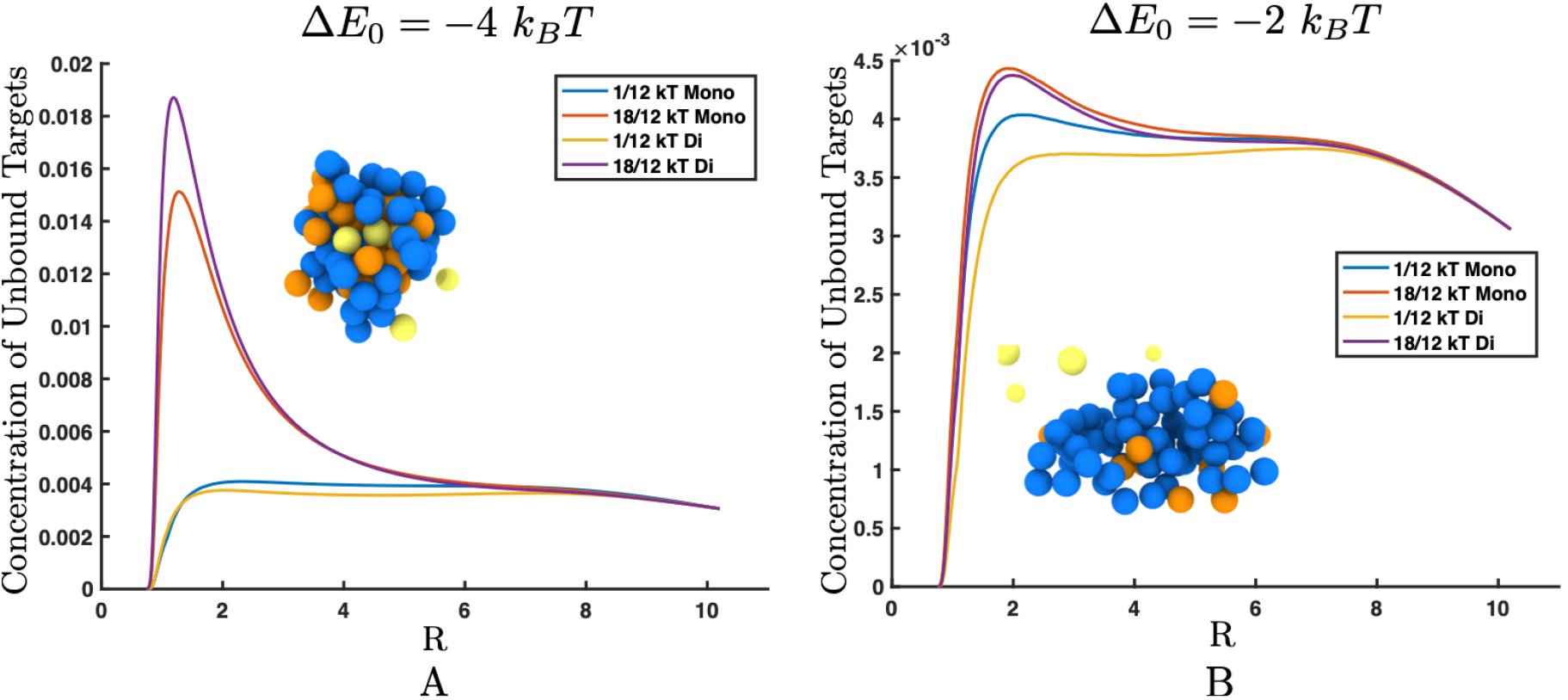
Plot of the minimum distance away from the polymer that unbound targets are found, normalized by the volume of a sphere with radius R, where R is the distance the center of the target is from the center of the nearest polymer bead. Data is shown for 64mer polymers in theta solvent with polymer-target binding affinities of (A) −4 *k*_*B*_*T* and (B) −2 *k*_*B*_*T*. (A) The concentration of unbound targets is approximately the same as the bulk when there is low inter-target attraction, but the concentration of unbound target near the polymer is higher than the bulk concentration when the inter-target potential is increased. The rendering in the inset shows unbound targets (yellow) clustered inside the polymer (blue) by the bound targets (orange). (B) Fewer targets have bound to the polymer, so the polymer has not collapsed. This makes the local concentration of unbound targets near the polymer approximately the same as the bulk concentration for both high and low inter-target attractions.

At a target-polymer binding affinity of −4*k*_*B*_*T*, this effect is not specific to the divalent targets, and increased aggregation can also be seen for monovalent targets, although less extreme. But at lower binding affinities such as −2*k*_*B*_*T* shown in Fig. 10B and Fig. 13B, targets are unaffected by target-target attraction because they do not bind strongly enough to the polymer to create the critical concentration needed on the polymer to attract more targets. Therefore, the average number of targets bound to the polymer barely increases at higher inter-target attraction.

Above a critical length or a critical binding affinity, polymers are able to take advantage of weakly attractive inter-target interactions, increasing inhibitor binding avidity. Though competition for binding sites lowers the *τ*_*B*_ for individual targets, inter-target attraction allows the polymer to induce a collapse transition that clusters unbound targets, significantly increasing the binding of the polymer overall. With a sharper collapse transition and a diminished entropy of collapse and re-swelling, longer polymers should show an amplified effect.

## 4 Conclusion

This work has shown that increasing the degree of polymerization of a multivalent inhibitor increases the overall avidity of binding, but there is a limited increase in avidity at high degrees of polymerization. To explore the effect of multivalent polymer structure, we used a Brownian dynamics bead-spring model coupled with a reactive polymer-target binding model to investigate how degree of polymerization influences a polymeric inhibitor’s avidity. First, we examined how the length of our inhibiting polymer modulates binding interactions with a single mono and divalent target model. We found that, consistent with previously reported experimental results for polymer binding to lectins, increasing the inhibitor length did increase binding avidity for multivalent targets, but interestingly, this effect was limited. We provide evidence that this limit can be explained by the entropic penalty of forming large loops; long polymers theoretically provide more possible loops when bound to a target in two places, but the entropic cost of forming long loops makes them unachievable in practice. Therefore, if the target is a protein, shorter polymers will create the maximum avidity. Due to its estimation of the targets as point particles, our model does not address the experimental results that increasing polymer length continues to increase avidity for larger many-valent targets such as viruses. Consequently, for targeting viruses, further investigation is necessary and researchers may want to continue creating polymers with higher degrees of polymerization.

In the presence of multiple targets, we found that longer polymers are able to use inter-target interactions to increase their avidity further. We show that despite decreased time bound for individual targets, longer polymers are able to bind to more targets simultaneously in the presence of favorable inter-target interactions. When inter-target attraction is present, longer polymers are able to induce a collapse transition where targets precipitate into a globule with the polymer, helping the polymer draw in a significant number of unbound targets. Increasing the concentration of unbound targets near the polymer makes the polymer better at clustering and binding targets. This could be a desirable effect in both the inhibition of targets and in other scenarios such as controlling biological signaling [34].

Our results suggest design rules for creating multivalent polymeric binders. With the understanding that increasing degree of polymerization has a limited effect on avidity in low target concentration environments and that inhibitor length can be used to induce phase separation in high concentration environments, future designers can focus on other variables when creating multivalent polymeric binders for proteins.

## 5 Author contributions

- Designed research (EZ, AAK)
- Performed research (EZ, JSW, AAK)
- Analyzed data (EZ, JSW, AAK)
- Wrote the manuscript (EZ, JSW, AAK)

## 6 Acknowledgements

The authors were supported by the Department of Defense (DoD) through the National Science and Engineering Graduate Fellowship (NDSEG) Program. The authors were also supported by the Ida M. Green Fellowship through the MIT Office of the Dean of Graduate Education. Computational resources were provided in part by the MIT Supercloud [35]. J. W. was supported by the National Science Foundation Graduate Research Fellowship under Grant No. 1122374. The authors would also like to thank Katharina Ribbeck and the Ribbeck Lab for many constructive discussions and suggestions.

## References

[1] Mathai Mammen, Seok-Ki Choi, and George M Whitesides. Polyvalent interactions in biological systems: implications for design and use of multivalent ligands and inhibitors. Angewandte Chemie, International Edition, 37(Copyright (C) 2016 American Chemical Society (ACS). All Rights Reserved.):2755–2794, 1998.

[2] Vijay M Krishnamurthy, Lara A. Estroff, and George M Whitesides. Multivalency in Ligand Design. In Raimund Mannhold, Hugo Kubinyi, Gerd Folkers, Wolfgang Jahnke, and Daniel A. Erlanson, editors, Fragment-based Approaches in Drug Discovery, chapter 2, pages 11–53. aug 2006.

[3] Laura L. Kiessling, Travis Young, and Kathleen H. Mortell. Multivalency in Protein-Carbohydrate Recognition. In Bertram O. Fraser-Reid, Kuniaki Tatsuta, and Joachim Thiem, editors, Glycoscience: Chemistry and Chemical Biology I-III, pages 1817–1861. Springer, Berlin, Heidelberg, 2001.

[4] Jeffrey D Esko and Nathan Sharon. Microbial Lectins: Hemagglutinins, Adhesins, and Toxins. In A Varki, R D Cummings, J D Esko, H H Freeze, P Stanley, C R Bertozzi, G W Hart, and M E Etzler, editors, Essentials of Glycobiology, chapter 34, pages 489–500. Cold Spring Harbor Laboratory Press, Cold Spring Harbor, NY, 2 edition, 2009.

[5] Sumati Bhatia, Mathias Dimde, and Rainer Haag. Multivalent glycoconjugates as vaccines and potential drug candidates. Med. Chem. Commun., 5(7):862–878, may 2014.

[6] Thomas R Branson and W Bruce Turnbull. Bacterial toxin inhibitors based on multivalent scaffolds. Chem. Soc. Rev., 42(11):4613–4622, 2013.

[7] David A. Rasko and Vanessa Sperandio. Anti-virulence strategies to combat bacteria-mediated disease. Nature Reviews Drug Discovery, 9(2)117–128, 2010.

[8] Shuang Liu and Kristi L Kiick. Architecture Effects on the Binding of Cholera Toxin by Helical Glycopolypeptides. Macromolecules, 41(3)764–772, feb 2008.

[9] David Deniaud, Karine Julienne, and Śebastien G Gouin. Insights in the rational design of synthetic multivalent glycoconjugates as lectin ligands. Org. Biomol. Chem., 9(4)966–979, 2011.

[10] Shengchang Tang, Wendy B. Puryear, Brian M. Seifried, Xuehui Dong, Jonathan A. Runstadler, Katharina Ribbeck, and Bradley D. Olsen. Antiviral Agents from Multivalent Presentation of Sialyl Oligosaccharides on Brush Polymers. ACS Macro Letters, 5(3)413–418, mar 2016.

[11] Ilona Papp, Christian Sieben, Adam L. Sisson, Johanna Kostka, Christoph Böttcher, Kai Ludwig, Andreas Herrmann, and Rainer Haag. Inhibition of Influenza Virus Activity by Multivalent Glycoarchitectures with Matched Sizes. ChemBioChem, 12(6)887–895, 2011.

[12] Joseph J Lundquist, Sheryl D Debenham, and Eric J Toone. Multivalency Effects in Protein-Carbohydrate Interaction: The Binding of the Shiga-like Toxin 1 Binding Subunit to Multi-valent C-Linked Glycopeptides. The Journal of Organic Chemistry, 65(24)8245–8250, dec 2000.

[13] Xiaoxiong Zeng, Takeomi Murata, Hirokazu Kawagishi, Taichi Usui, and Kazukiyo Kobayashi. Synthesis of Artificial N-Glycopolypeptides Carrying N-Acetyllactosamine and Related Compounds and Their Specific Interactions with Lectins. Bioscience, Biotechnology, and Biochemistry, 62(6)1171–1178, jan 1998.

[14] Miho Watanabe, Koji Matsuoka, Eiji Kita, Katsura Igai, Nobutaka Higashi, Atsushi Miyagawa, Toshiyuki Watanabe, Ryohei Yanoshita, Yuji Samejima, Daiyo Terunuma, Yasuhiro Natori, and Kiyotaka Nishikawa. Oral Therapeutic Agents with Highly Clustered Globotriose for Treatment of Shiga Toxigenic Escherichia coli Infections. The Journal of Infectious Diseases, 189(3)360–368, 2004.

[15] Brian D Polizzotti and Kristi L Kiick. Effects of Polymer Structure on the Inhibition of Cholera Toxin by Linear Polypeptide-Based Glycopolymers. Biomacromolecules, 7(2)483–490, feb 2006.

[16] Brian D Polizzotti, Ronak Maheshwari, Jan Vinkenborg, and Kristi L Kiick. Effects of Saccharide Spacing and Chain Extension on Toxin Inhibition by Glycopolypeptides of Well-Defined Architecture. Macromolecules, 40(20)7103–7110, oct 2007.

[17] Susanne Liese and Roland R. Netz. Influence of length and flexibility of spacers on the binding affinity of divalent ligands. Beilstein Journal of Organic Chemistry, 11:804–816, 2015.

[18] George B Sigal, Mathai Mammen, Georg Dahmann, and George M Whitesides. Polyacrylamides Bearing Pendant α-Sialoside Groups Strongly Inhibit Agglutination of Erythrocytes by Influenza Virus: The Strong Inhibition Reflects Enhanced Binding through Cooperative Polyvalent Interactions. Journal of the American Chemical Society, 118(16)3789–3800, jan 1996.

[19] Masanori Nagao, Yurina Fujiwara, Teruhiko Matsubara, Yu Hoshino, Toshinori Sato, and Yoshiko Miura. Design of Glycopolymers Carrying Sialyl Oligosaccharides for Controlling the Interaction with the Influenza Virus. Biomacromolecules, 18(12)4385–4392, dec 2017.

[20] Motomu Kanai, Kathleen H Mortell, and Laura L Kiessling. Varying the Size of Multivalent Ligands: The Dependence of Concanavalin A Binding on Neoglycopolymer Length. Journal of the American Chemical Society, 119(41)9931–9932, oct 1997.

[21] Sarah-Jane Richards, Mathew W. Jones, Mark Hunaban, David M. Haddleton, and Matthew I. Gibson. Probing Bacterial-Toxin Inhibition with Synthetic Glycopolymers Prepared by Tandem Post-Polymerization Modification: Role of Linker Length and Carbohydrate Density. Angewandte Chemie International Edition, 51(31)7812–7816, jul 2012.

[22] Helen M Berman, John Westbrook, Zukang Feng, Gary Gilliland, T N Bhat, Helge Weissig, Ilya N. Shindyalov, and Philip E. Bourne. The Protein Data Bank. Nucleic Acids Research, 28(1)235–242, jan 2000.

[23] R. Ravishankar, C.J. Thomas, K. Suguna, A. Surolia, and M. Vijayan. Crystal structures of the peanut lectin-lactose complex at acidic pH: retention of unusual quaternary structure, empty and carbohydrate bound combining sites, molecular mimicry and crystal packing directed by inte. Proteins, 43:260–270, 2001.

[24] Alfredo Alexander-Katz and Roland R Netz. Dynamics and Instabilities of Collapsed Polymers in Shear Flow. Macromolecules, 41(9)3363–3374, may 2008.

[25] Charles E Sing and Alfredo Alexander-Katz. Equilibrium Structure and Dynamics of Self-Associating Single Polymers. Macromolecules, 44(17)6962–6971, sep 2011.

[26] Moira Ambrosi, Neil R Cameron, Benjamin G Davis, and Snjezana Stolnik. Investigation of the interaction between peanut agglutinin and synthetic glycopolymeric multivalent ligands. Organic & Biomolecular Chemistry, 3(8)1476, 2005.

[27] René Roy. Syntheses and some applications of chemically defined multivalent glycoconjugates. Current Opinion in Structural Biology, 6(5)692–702, oct 1996.

[28] D J Diestler and E W Knapp. Statistical Mechanics of the Stability of Multivalent Ligand-Receptor Complexes †. The Journal of Physical Chemistry C, 114(12)5287–5304, apr 2010.

[29] William P Jencks. On the attribution and additivity of binding energies (proteins/ligands/entropy/enzymes). Biochemistry, 78(7)4046–4050, 1981.

[30] Marcus Weber, Alexander Bujotzek, and Rainer Haag. Quantifying the rebinding effect in multivalent chemical ligand-receptor systems. Journal of Chemical Physics, 137(5)1–11, 2012.

[31] Douglas Poland and Harold A Scheraga. Phase Transitions in One Dimension and the Helix-Coil Transition in Polyamino Acids. The Journal of Chemical Physics, 45(5)1456–1463, sep 1966.

[32] Dreania Levine, Michael J Kaplan, and Peter J Greenaway. The Purification and Characterization of Wheat-Germ Agglutinin. Technical report, 1972.

[33] 5 Human Albumin. Transfusion Medicine and Hemotherapy, 36(6)399–407, 2009.

[34] Christopher W Cairo, Jason E Gestwicki, Motomu Kanai, and Laura L Kiessling. Control of Multivalent Interactions by Binding Epitope Density. Journal of the American Chemical Society, 124(8)1615–1619, feb 2002.

[35] Albert Reuther, Jeremy Kepner, William Arcand, David Bestor, Bill Bergeron, Chansup Byun, Matthew Hubbell, Peter Michaleas, Julie Mullen, Andrew Prout, and Antonio Rosa. LLSuper-Cloud: Sharing HPC systems for diverse rapid prototyping. In 2013 IEEE High Performance Extreme Computing Conference (HPEC), pages 1–6. IEEE, sep 2013.

